# Genomic evidence of *Escherichia coli* gut population diversity translocation in leukemia patients

**DOI:** 10.1101/2024.06.25.600660

**Authors:** Julie Marin, Violaine Walewski, Thorsten Braun, Samira Dziri, Mélanie Magnan, Erick Denamur, Etienne Carbonnelle, Antoine Bridier-Nahmias

## Abstract

*Escherichia coli*, a commensal species of the human gut, is an opportunistic pathogen which can reach extra-intestinal compartments, including the bloodstream and the bladder, among others. In non-immunosuppressed patients, purifying or neutral evolution of *E. coli* populations has been reported in the gut. Conversely, it has been suggested that when migrating to extra-intestinal compartments, *E. coli* genomes undergo diversifying selection as supported by strong evidence for adaptation. The level of genomic polymorphism and the size of the populations translocating from gut to extra-intestinal compartments is largely unknown.

To gain insights in the pathophysiology of these translocations, we investigated the level of polymorphism and the evolutionary forces acting on the genomes of 77 *E. coli* isolated from various compartments in three immunosuppressed patients. Each patient had a unique strain which was a mutator in one case. In all instances, we observed that translocation encompasses the majority of the genomic diversity present in the gut. The same signature of selection, whether purifying or diversifying, and as anticipated, neutral for mutator isolates, was observed in both the gut and bloodstream. Additionally, we found a limited number of non-specific mutations among compartments for non-mutator isolates. In all cases, urine isolates were dominated by neutral selection. These findings indicate that substantial proportions of populations are undergoing translocation and that they present a complex compartment-specific pattern of selection at the patient level.

**Importance:** It has been suggested that intra and extra-intestinal compartments differentially constrain the evolution of *E. coli* strains. Whether host particular conditions, such as immunosuppression, could affect the strain evolutionary trajectories remain understudied. We found that, in immunosuppressed patients, large fractions of *E. coli* gut populations are translocating with variable modifications of the signature of selection for commensal and pathogenic isolates according to the compartment and/or the patient. Such multiple site sampling should be performed in large cohorts of patients to get a better understanding of *E. coli* extra-intestinal diseases.

## Introduction

Bloodstream infections (BSIs) are still a major concern among onco-hematologic patients, influenced by factors such as the type of pathogen, the degree of host immunodeficiency and the status of underlying disease. Despite advances in the clinical management of hematological malignancies, BSIs remain life-threatening complications in the clinical course of these patients, with reported crude mortality rate up to 40% (1–3). A clear shift in the bacterial species causing BSIs in patients with hematological malignancies has been recently reported transitioning from Gram-positive to Gram-negative, in the first place Enterobacteriaceae, and in particular *Escherichia coli*, represent the most frequently involved bacterial species, together with the worrisome and growing phenomenon of multiresistant bacteria (4, 5).

*E. coli* is a commensal species of the lower intestine of humans (6). The gut is its primary habitat, and probably the main ecological context of selection. Virulence genes, for instance, are thought to be primarily selected in the intestine as a by-product of commensalism (7, 8). *E. coli* is also an opportunistic pathogen, frequently responsible for intestinal and extra-intestinal infections (9). When reaching a new compartment, such as the bladder or the bloodstream, *E. coli* faces new challenges and new opportunities for adaptation.

Previous studies have mainly investigated the signatures of selection in commensal *E. coli* isolates and those sampled from extra-intestinal infections, revealing different scenarios. The evolution of commensal *E. coli* has been found to be governed by purifying selection, whether it implies the entire genome (10) or specific genes such as the H7 flagellin genes (11). However, another study, following the evolution of a clone in a single individual, did not find any evidence for selection in the gut (12). On the contrary, adaptation during chronic and acute infections has been observed (13). In particular, it has been shown that *E. coli* isolates colonizing extra-intestinal sites were adapting under strong selective pressure, with an excess of non-synonymous mutations and patterns of convergence at the gene level (including for H7 flagellin genes). Evidence for adaptation during chronic infection have been emphasized for other bacteria, such as *Burkholderia dolosa* (14) or *Pseudomonas aeruginosa* (15–17), with genotypically and phenotypically diversifying lineages during cystic fibrosis infections for instance (18–21). Adaptation to the human host (22, 23), evasion from the immune response (24, 25) and acquisition of antibiotic resistances (26, 27) are favored by this accumulation of mutations. In addition, niche adaptation shapes the allelic diversity among compartments with specific mutations occurring in the gut or in the bladder associated with functions increasing *E. coli* fitness in each respective compartment (28).

What happens when the intestinal barrier and the immune response are weakened? Patient deficiencies, such as immunosuppression, weakening of the intestinal barrier by antibiotic therapy or chemotherapy, can increased the risk of *E. coli* extra-intestinal infections (29–31). Anticancer chemotherapy drugs directly impact the intestinal microbiota (32), leading to dysbiosis and subsequent intestinal mucositis which increase the risk of bacterial translocation to the bloodstream (33). Moreover, among patients undergoing chemotherapy, those with leukemia are highly prone to extra-intestinal infections and relapse (34). The weakening of intestinal barriers and the lack of immune defense could alter the adaptive conditions described for commensal and pathogenic isolates of *E. coli* in non-immunosuppressed patients. Modifications of the signature of selection are therefore expected for commensal and pathogenic isolates of *E. coli* in immunosuppressed patients. In addition, the level of genomic polymorphism and the proportion of the population translocating is largely unknown.

Here, we evaluated the selective forces acting on *E. coli* evolution in three immunosuppressed patients among three compartments: bladder, bloodstream and gut. For each patient, we analyzed whole genome sequences from isolates found in each compartment to assess the genetic diversity and the strength of the various selective processes at play.

## Material and methods

### Sampling

Clinical isolates were isolated from three patients with leukemia and *E. coli* sepsis hospitalized in Avicenne hospital (Seine-Saint-Denis, France). Patient A experienced three infectious episodes while patient B and C had one each. For each episode, these patients had at least one blood culture positive for *E. coli*. Isolates were sampled from positive blood culture and from bladder and feces samples on the same or subsequent day. The initial blood culture was obtained before any antibiotic therapy in patients newly admitted to the hospital. All procedures performed were in accordance with the ethical standards of the responsible committee on human experimentation (institutional and national), validated by the ethics committee of Avicenne hospital (Comité Local d’Ethique d’Avicenne). Information on the type and status of hematologic disease, presence of neutropenia (<0.5×10^9^/L), previous exposure to any antibiotic therapy, including prophylaxis or treatment of prior infectious episodes, type of infection, microbiological isolate, and outcome was collected in a database.

### Patient characteristics

The three patients (A to C) studied had hematological pathologies: two with acute myeloid leukemia and one with Hodgkin’s lymphoma. Patient A was diagnosed with acute myeloid leukemia and myelofibrosis in January 2014 (Acute myeloblastic leukemia with minimal maturation M1), treated with an allograft. He experienced relapse in March 2015, with febrile neutropenia and presence of circulating blasts. During the BSI, the patient presented an inflammatory syndrome, a severe immunosuppression and had received antibiotics in the previous 30 days. Patient B suffered from undifferentiated acute myeloblastic leukemia (M0) diagnosed and allografted in 2015. At the time of BSI, he was receiving chemotherapy, had severe immunosuppression, an inflammatory syndrome, and had received antibiotics within the preceding 30 days. Patient C was diagnosed with Hodgkin’s lymphoma in 2009. Relapsed in 2017 during the BSI episode, the patient exhibited inflammatory syndrome, severe immunosuppression and had previously received antibiotics.

### Prior in vitro phylogroup typing

Phylogroup of each sample (blood, urinary or feces) were determined using quadruplex PCR (35) to select isolates belonging to the same phylogroup for each infection episode.

### Genome sequencing and assembling

Whole-genome sequencing was performed on each sample using Illumina Technology (MiSeq and HiSeq 2500) and Nextera XT library preparation kits as instructed by the manufacturer (Illumina, San Diego, CA). Fastq files (raw sequencing data) were submitted to the European nucleotide archive (see **Table S1** for accession numbers). Genome assembly was performed with SPAdes v.3.15.5 (36) (see **Table S1** for assembly quality results).

For each patient, one strain was chosen as a reference and was sequenced with the Oxford Nanopore Technologies MinION platform using an R9.4 flow cell. We prepared the samples with kits LSK-108 and NBD-104 for library preparation and barcoding. The Nanopore reads were filtered with Filtlong (37) using the following parameters: minimum length of 1000 and 95% of the best reads kept. High-quality assemblies of the three reference genomes were assembled with a hybrid strategy, using both Illumina and Nanopore reads with Unicycler V0.4.4 (38). Information on the hybrid assembly quality is presented in **Table S2**.

### Core and accessory genome

Seventy-seven SPAdes assemblies were annotated with Prokka (39). Plasmid sequences were predicted by PlaScope (40). Pan-genome analysis from annotated assemblies were performed with Roary using default parameters (41). The core genome alignment and the list of genes of the accessory genome were generated for the 3 patients.

### Variant calling (SNPs and deletions)

Bases with a low-quality score (< 30) were discarded and the adapters were removed with Trim Galore [a wrapper of the Cutadapt program (42)]. SNPs were detected by aligning the reads to the corresponding reference sequence of each patient (**Table S1**) with Snippy 4.4.0 (43) with the following parameters: the nucleotide minimum quality to be analyzed (basequal) equal to 20, the minimum number of reads covering a site (mincov) equal to 10 and the minimum proportion of those reads different from the reference (minfrac) equal to 0.9. Structural variants (deletions) were detected by mapping reads to the corresponding reference assembly with BWA-MEM (44) and then Sequence Alignment Map (sam) files were analyzed with Wham (45). As recommended, we remove calls smaller than 50 bp and larger than 2 Mbs. For each strain, we also removed calls with less than 5 supporting reads.

### Detection of insertion sequence (IS) elements

IS elements on the three reference sequences were identified with ISFinder (46). We selected hits with an e-value lower than 10e-10, a minimum alignment coverage of 50% and a minimum identity of 70%. Next, we searched for differences in the IS repertoire for each isolate against the corresponding reference with panISa v0.1.6 (47).

### Typing and genotypic antibiotic resistance and virulence

We used an in-house script, Petanc (48), that integrates several existing bacterial genomic tools to perform the typing of isolates with several genotyping schemes using the genomic tool SRST2 (49). Sequence types (STs) were defined using the Warwick MLST scheme and the Pasteur scheme (50, 51). We only used the Warwick scheme for the analyses described hereafter. We also determined the O:H serotypes and the FimH alleles (52, 53). The phylogroups were confirmed using the ClermonTyping method (54).

The resistome and virulome were first established using the in-house script Petanc (48). They were defined by BlastN with Abricate (https://github.com/tseemann/abricate) using the ResFinder (version 4.2.2) database (55), a custom database including the VirulenceFinder database and VFDB (56, 57), to which we added selected genes (58). We set the threshold for minimum identity to 80% with a minimum coverage of 90%. Next, a pan-resistome and pan-virulome were built including all the antibiotic resistance and virulence associated genes of all isolates. We mapped the reads to the corresponding pan-resistome and pan-virulome with BWA-MEM (44). We considered a gene as present when we found more than 80% coverage and at least one read. We also tested more conservative thresholds with 80% coverage and more than 5 reads.

To evaluate the possibility of false-negative results, we applied the same methodology for 14 MLST genes (*adk*, *dinB*, *fumC*, *gyrB*, *icd*, *mdh*, *pabB*, *polB*, *purA*, *putP*, *recA*, *trpA*, *trpB*, *uidA*) and for genes encoded on a plasmid (predicted by PlaScope (40)), 157, 381 and 157 genes for patient A, B, and C, respectively).

### Genomic diversity and traces of selection

Rates of nonsynonymous and synonymous mutation were compared by computing non-synonymous substitution / synonymous substitution (dN/dS) ratios (R language (59)) from the gene alignments obtained with Roary, to evaluate the genomic traces of selection. To be able to compare the isolates taking into account their phylogenetic history, we reconstructed an ancestral sequence for each patient to which each isolate was then compared. We first midpoint rooted the trees of patient B and C for which there was only one sampling time. For patient A, we rooted the tree based on the best root-to-tip correlation (function ‘initRoot’, package BactDating (60)). Next we inferred the ancestral sequence as the sequence at the root of each tree using parsimony (function ancestral.pars, package phangorn (61)). We calculated dN and dS as the observed number of substitutions of each type divided by the number of potential substitutions of the same type in the considered sequence. For each codon, the number of potential non-synonymous or synonymous substitutions is determined by the genetic code. We then computed the dN/dS ratio for each pair of sequences, we determined the mean dN/dS ratio and assessed whether it significantly differed from 1 (Wilcoxon test), where 1 represents perfect balance between diversifying and purifying selection indicating no visible selection. When in a pair of sequences, one or more non-synonymous and zero synonymous mutations are encountered, an infinite value is returned. To take into account these non-synonymous mutations, we used the Laplace smoothing technique (62), which is used to overcome issues caused by certain values having zero occurrence. We computed the standard deviation of all possible synonymous mutations and added this value to each term of the dN/dS ratio. When there was no mutation, neither synonymous nor non-synonymous, we removed this comparison because this case did not correspond to any of the three categories, diversifying, purifying or neutral selection.

To examine changes that occurred specifically in each compartment, we used the same methodology and computed dN/dS ratios for clades grouping samples of the same compartment. Here, the ancestral sequence was defined as the inferred sequence at the root of the focal clade.

## Results

We evaluated the genomic diversity (SNPs and deletions) and the genomic traces of selection in 77 *E. coli* isolates sampled concomitantly from the gut, blood and urine compartments of three patients (**Figure 1**). The distribution of isolates for each patient and compartment is detailed in **Table 1**.

**Figure 1.**
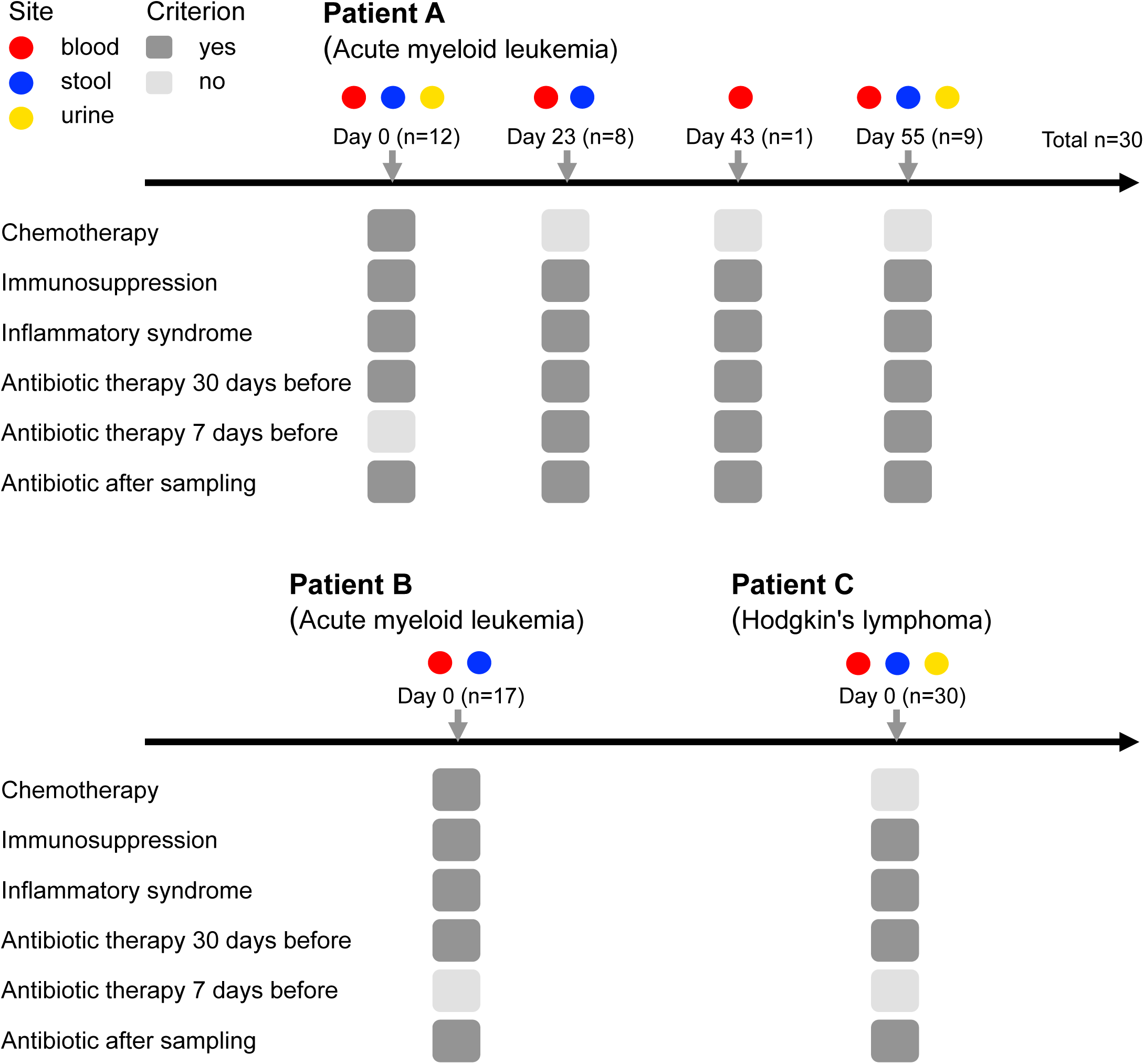
Patient follow-up with their main clinical characteristics and the sampling schemes. Absence (light gray) or presence (dark grey) of each criterion are shown. Patient A had four infectious episodes, patients B and C had one infectious episode.

**Table 1.** Patient sampling and strain typing.

| Patient | Number of isolates<br>(stool – blood – urine) † | Phylogroup | Sequence type | Serotype | *fimH* allele |
| --- | --- | --- | --- | --- | --- |
| Patient A | 30 (13 -12 - 5) | A | 10 | O101:H9 | *fimH54* |
| Patient B* | 17 (10 - 7) | B1 | 1431 | O8:H19 | *fimH32* |
| Patient C | 30 (10 - 10 - 10) | B2 | 131 | O25b:H4 | *fimH30* |

Phylogroups, MLSTs (Warwick and Pasteur), serotypes and *fimH* alleles are indicated. All the isolates of patient B were mutators (\*).
(+) Patients B and C were sampled at a single time point whereas samples of patient A correspond to 4 timepoints.

### Phylogroup, ST, serotype and fimH allele diversity

Within each infection episode, all isolates belonged to the same phylogroup and ST and had the same serotype and *fimH* allele (**Table 1**). Isolates belonged to the phylogroup A (patient A), B1 (patients B) and B2 (patient C).

### Global genomic diversity (SNPs and IS)

We first looked at the SNP diversity. With the exception of patient B, isolates exhibited a low number of mutated genes (between 8 and 11) and no deletion (**Table S3**). Patient B isolates however, revealed 328 mutated genes. Indeed, we noted a deletion of *mutS* in patient B isolates. This inactivation of the DNA mismatch repair system conferred to them a mutator phenotype (63, 64) (**Table S4**).

In line with those results, we found a low number of SNPs in isolates from patients A and C (mean number of 5.07 [95% confidence interval: 4.20-5.95]) compared to those of patient B (mean number of 117.13 [82.16-152.10], p-value = 1.237e-08 (Wilcoxon test)) (**Figure 2**). Similarly, intra- and extra-intestinal isolates did not differ significantly in term of variant categories (**Table 2**) (chi-squared test after correcting for sample size, p-value = 0.99). No common genes with SNPs (**Table S5)** were identified among patients (**Table S3**).

**Figure 2.**
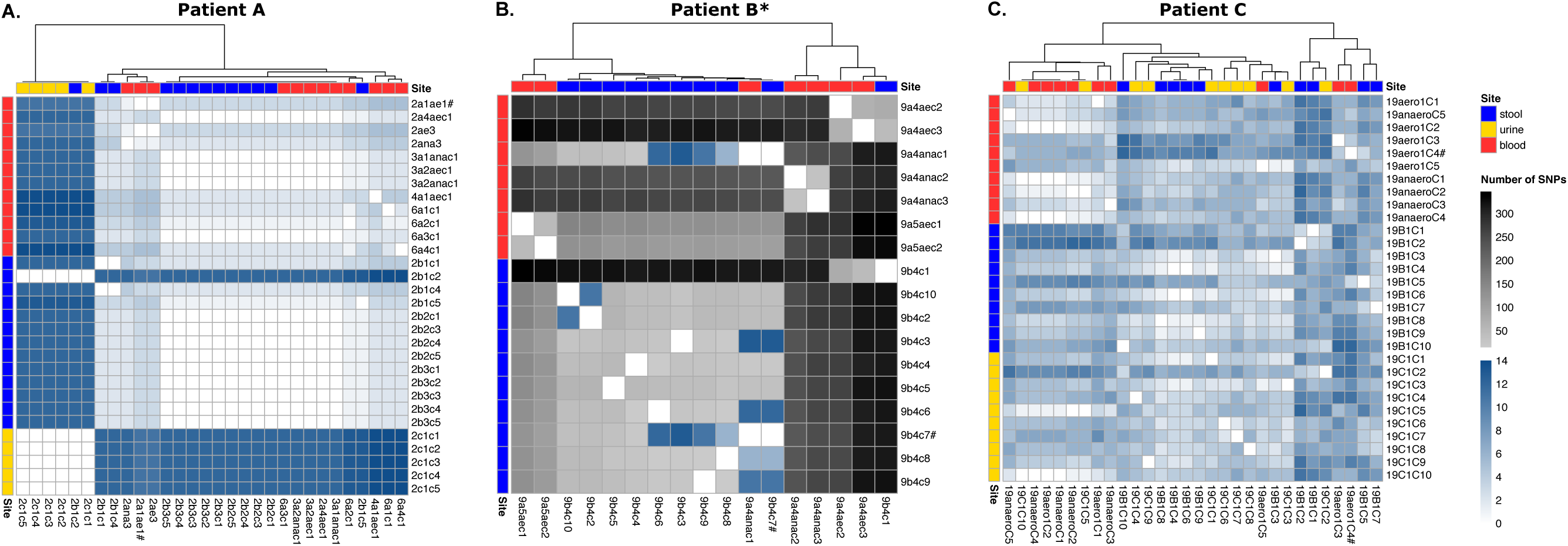
Genomic diversity of *E. coli* isolates among samples and compartments for the three patients. (A-C) Heatmaps showing the number of SNPs between each isolate of patients (A, B and C). Samples are ordered by site horizontally and clustered according to their SNP similarities vertically (method: complete). All isolates of patient B are mutators (*). The isolate corresponding to the reference sequence is indicated (#). We used the same color scale for all patients. Note that the number of SNPs for patients A and C is between 0 and 14 and between 0 and 400 for patient B.

**Table 2.** Number of variant types among isolates of a patient’s infection episode found with snippy.

|  | Site | conservative<br>inframe<br>insertion | disruptive<br>inframe<br>deletion | frameshift<br>variant | missense<br>variant | non coding<br>transcript<br>variant | stop<br>gained | stop lost | synonymous<br>variant |
| --- | --- | --- | --- | --- | --- | --- | --- | --- | --- |
| Patient A | blood<br>(n=12) | 9 | 0 | 1 | 3 | 0 | 1 | 0 | 9 |
|  | stool<br>(n = 13) | 12 | 1 | 0 | 6 | 0 | 0 | 1 | 17 |
|  | urine<br>(n=5) | 0 | 5 | 0 | 20 | 0 | 0 | 5 | 20 |
| Patient B* | blood<br>(n=7) | 0 | 0 | 216 | 541 | 5 | 11 | 20 | 292 |
|  | stool<br>(n=10) | 0 | 0 | 86 | 177 | 1 | 4 | 4 | 146 |
| Patient C | blood<br>(n=10) | 0 | 0 | 9 | 7 | 0 | 0 | 0 | 2 |
|  | stool<br>(n=10) | 0 | 0 | 16 | 26 | 0 | 0 | 0 | 14 |
|  | urine<br>(n=10) | 0 | 0 | 12 | 20 | 0 | 0 | 0 | 6 |
| Intra-intestinal<br>isolates | stool<br>(n=23) | 12 | 1 | 16 | 32 | 0 | 0 | 1 | 31 |
| Extra-intestinal<br>isolates | blood/urine<br>(n=37) | 9 | 5 | 22 | 50 | 0 | 1 | 5 | 37 |

We excluded patient B from intra- and extra-intestinal isolates because all the 17 isolates were
mutators (\*). n: number of isolates.

Additionally, the diversity in IS elements revealed only few differences among isolates of the same patient (0 to 5 for non-mutator strains) (**Tables S6-S7**). The number of IS within the isolates of each patient was in the upper part of the range found for *E. coli* genomes (65), with a lower number of IS elements for the mutator strain, as expected (66). We detected 5 supplementary potential IS elements in 1 to 14 isolates (4.6 isolates in average) for patient A compared to the reference genome (189 IS elements). For patient B (mutator strain), we detected 22 supplementary potential IS elements in 1 to 11 isolates (2.63 isolates in average) compared to the reference genome (97 IS elements). We did not find any supplementary potential IS element in patient C isolates compared to the reference genome (136 IS elements).

### Within patient compartment diversity

All isolates from a given patient were highly similar (except for patient B with mutator isolates) (**Figures 2-3**). Consequently, no large deletions were detected because we used one of these isolates as the reference strain for each patient (**Table S3**).

**Figure 3.**
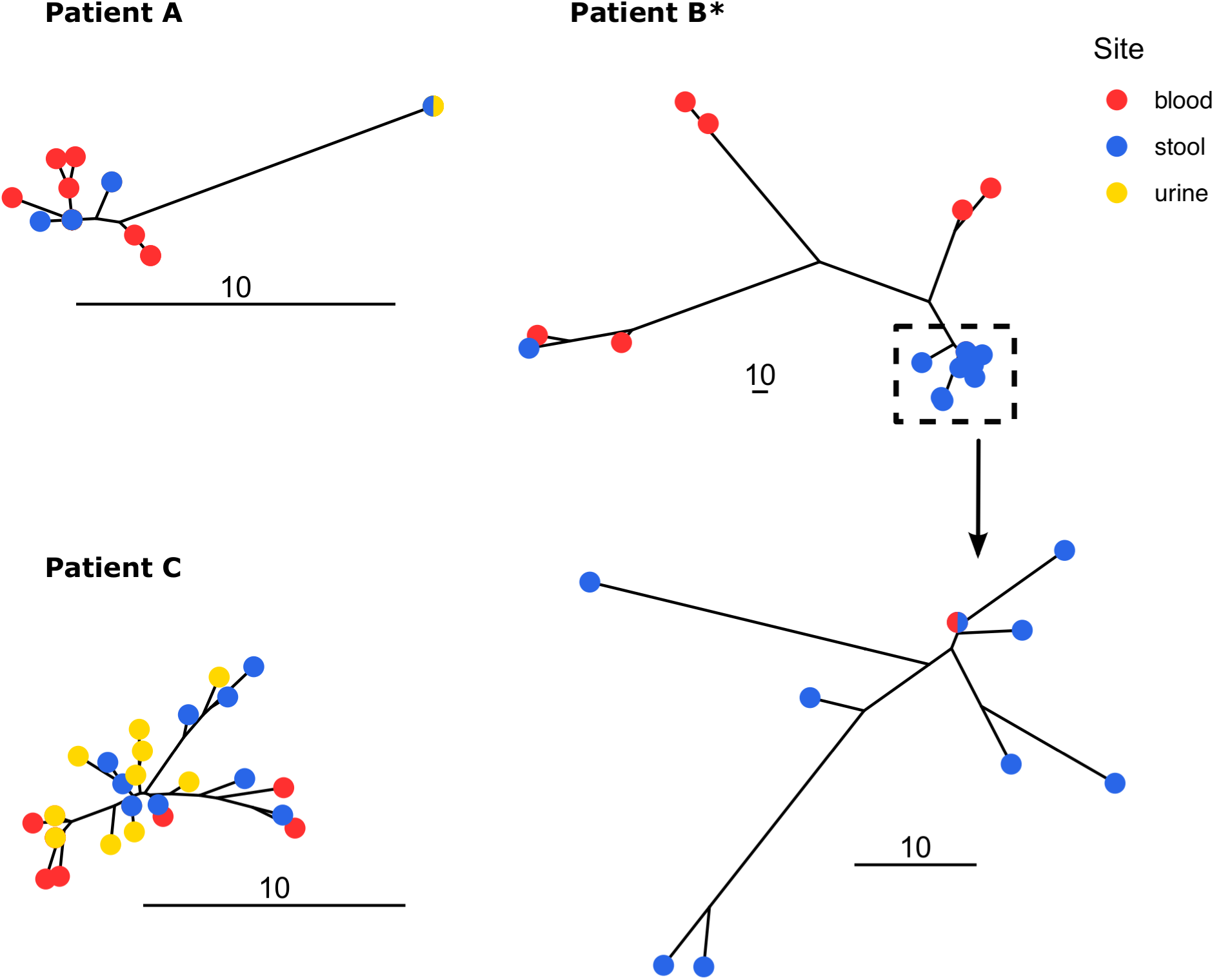
Unrooted trees of *E. coli* isolates from patients A, B and C. Trees were built using neighbor joining from the substitution presence/absence matrix. The scale indicates the number of substitutions. All isolates of patient B are mutators (*). We zoomed in on a clade of the patient B to highlight the scale difference. The bicolor points (patients A and C) denote the presence of isolates sampled in different sites with identical sequences (0 SNP).

The majority of SNPs were shared among compartments (**Figure 4**). For patient A, 13 over 20 SNPs were identical among at least 2 compartments and 13 over 22 for patient C. For patient B (mutator strain), despite a high level of polymorphism, most SNPs were shared among stool and blood isolates. Furthermore, most of the stool polymorphism was found in other compartments, 87%, 77% and 87% for patient A, B and C respectively, suggesting a large population translocation from the gut.

**Figure 4.**
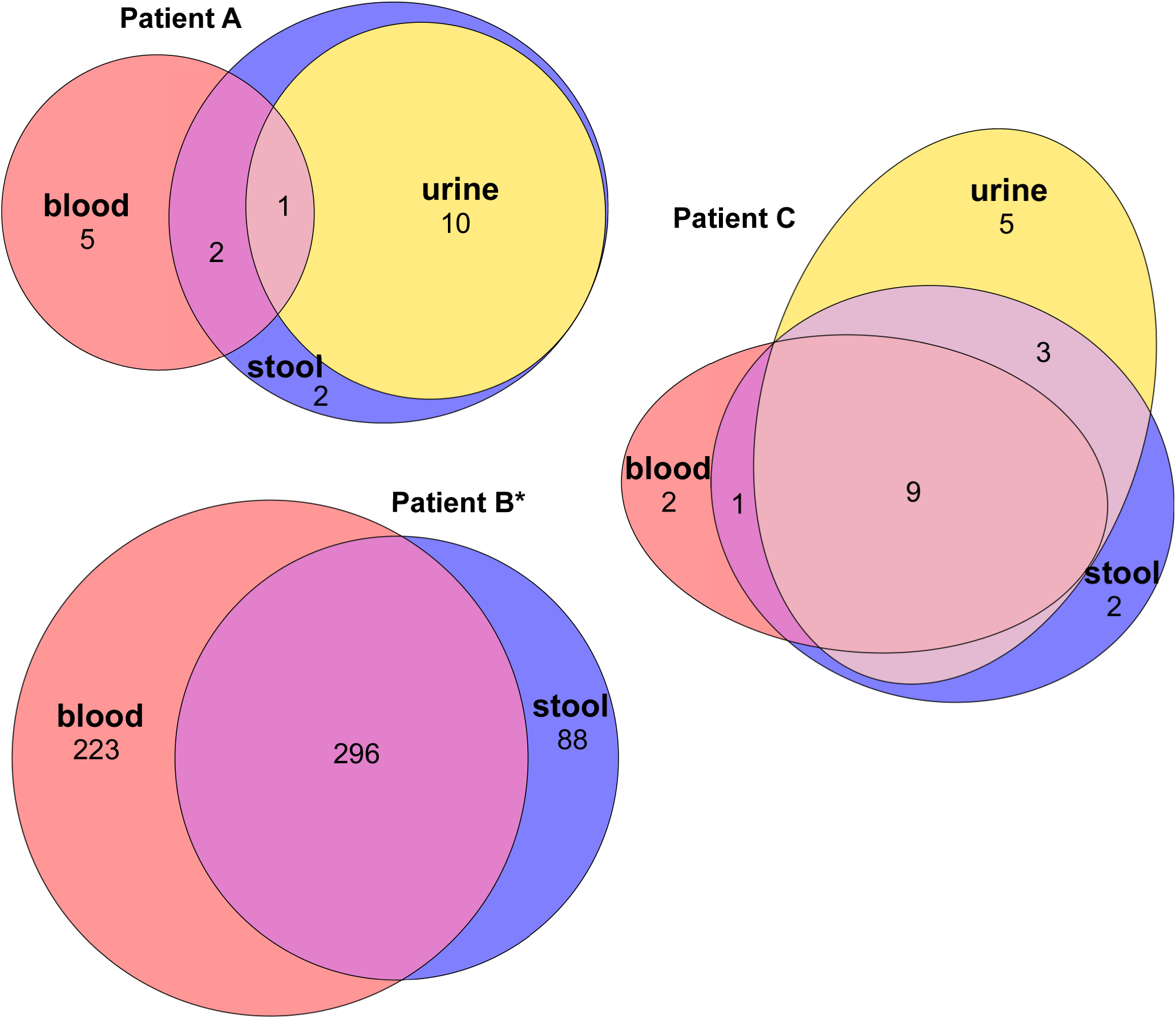
Venn diagrams showing the SNPs distribution among compartments for patients A, B and C. The ellipses are proportional to the number of SNPs. All isolates of patient B are mutators (*).

Additional IS elements were also shared among compartments (**Table S7**). For patient A, 3 over 5 additional IS elements were shared among 2 compartments and 12 over 22 for patient B.

### Antibiotic resistance and virulence associated gene content

Regarding genes associated with antibiotic resistance, isolates within each infection were highly similar. Limited variations were observed among isolates of patient B (mutator isolates) and A which were sampled at different time points (**Figures 1 and 5**). For patient A, we found gene presence/absence discrepancies for three resistance genes, *sul2*, *tet(A)* and *dfrA5*. The gene *sul2* was consistently predicted as encoded on a plasmid across all isolates (**Table S8**). The genes *dfrA5* and *tet(A)* were found on the same contig in 20 isolates out of 23 isolates possessing both genes, and were predicted as encoded on a plasmid for 96.67% and 86.96% of the isolates respectively. For patient B, we found presence/absence discrepancies for all the 7 resistance genes detected. The genes *qnrS1*, *bla*CTX-M-15, *bla*TEM-1B, *aph*(6)-Id, *aph*(3’’)-Ib and *sul2* were always co-located on the same contig and predicted as encoded on a plasmid. The gene *dfrA*14 was predicted as encoded on a plasmid for all isolates. Similar results were obtained using a more conservative threshold (at least 5 reads covering more than 80% of the gene) (**Figure S1**), with the exception of the gene *mdf(A)*, predicted as chromosomal, which was missing in three isolates of patient B.

**Figure 5.**
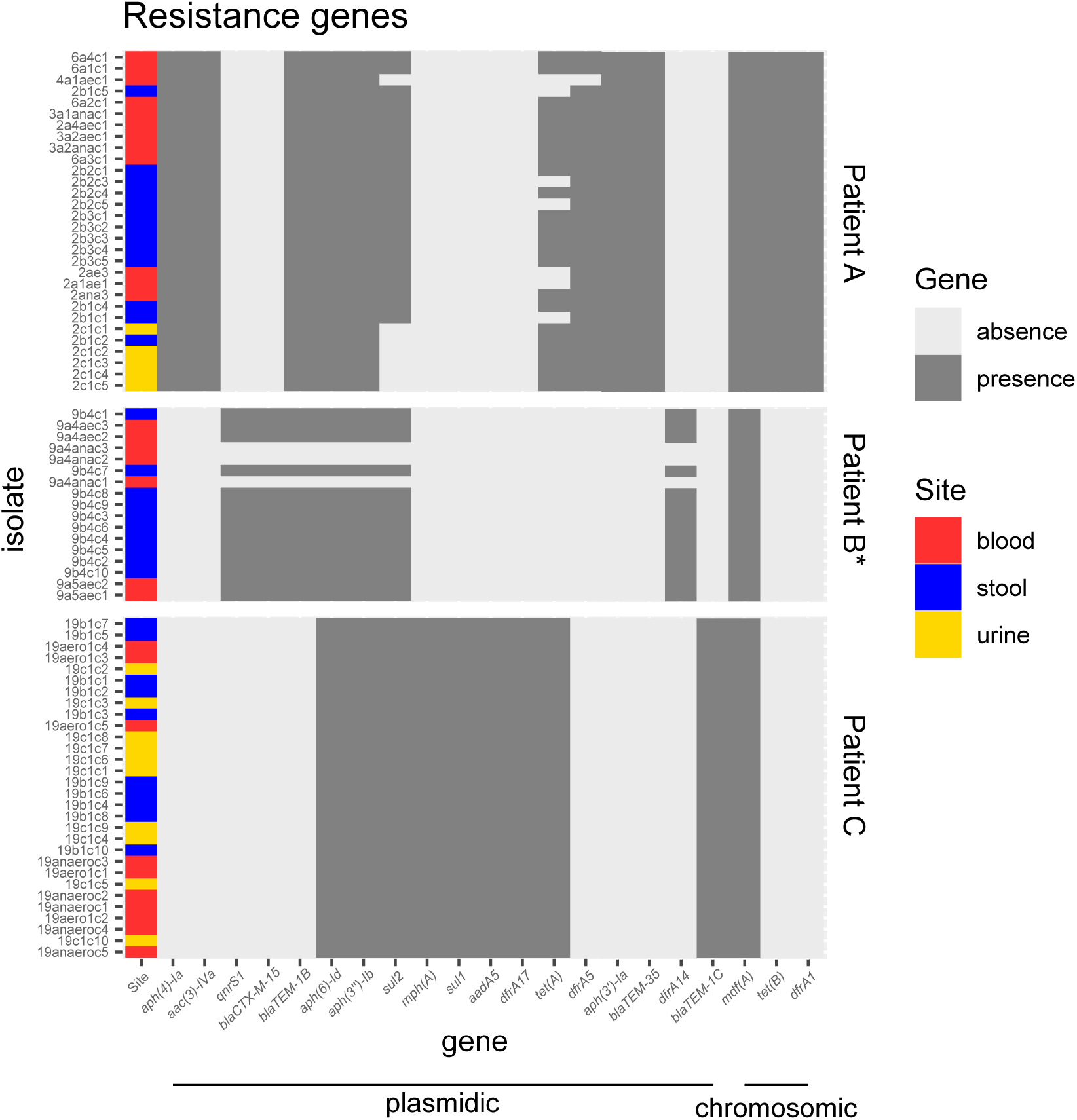
Presence/absence heatmaps of antibiotic resistance genes of *E. coli* isolates when compared to the pan-resistome (including the resistance genes of all isolates). We considered a gene as present when at least 80% of its length was covered by at least one read. Genes are ordered by synteny on contigs. All isolates of patient B are mutators (*). The prevailing predicted localization of genes by PlaScope (chromosomic or plasmidic) is indicated (full list in supplementary Table S8). Note that chromosomal genes are not mobile.

High gene content similarity was also observed for virulence associated genes (**Figure 6**). An identical gene content was found in patient A isolates. For patient C, we noted a discrepancy for a single gene, *iss*11, consistently predicted as chromosomal, and absent in four isolates. Slightly more differences were found in patient B isolates (mutator strain). The genes *espX5* and *espX1* were missing for one isolate and *gad*20 for five isolates, all predicted as chromosomal (**Table S9**). We found more discrepancies with a more conservative threshold (at least 5 reads covering more than 80% of the gene), highlighting lower sequencing depth of two isolates in particular (**Figure S2**). In patient A, all the isolates had the same gene content as the exception of 2 isolates (2ana3 and 9b4c8). We found six missing genes for 2ana3 and one missing gene for 2c1c5, all predicted as chromosomal. In addition, *entA*, *entB* genes were always adjacent on the same contig (**Table S9**). In patient C isolates, the gene *iss*11, always predicted as chromosomal, was missing in 5 isolates. Patient B displayed a greater number of differences, over the 47 virulence-associated genes detected, all predicted as chromosomal, only 15 were present in all isolates.

**Figure 6.**
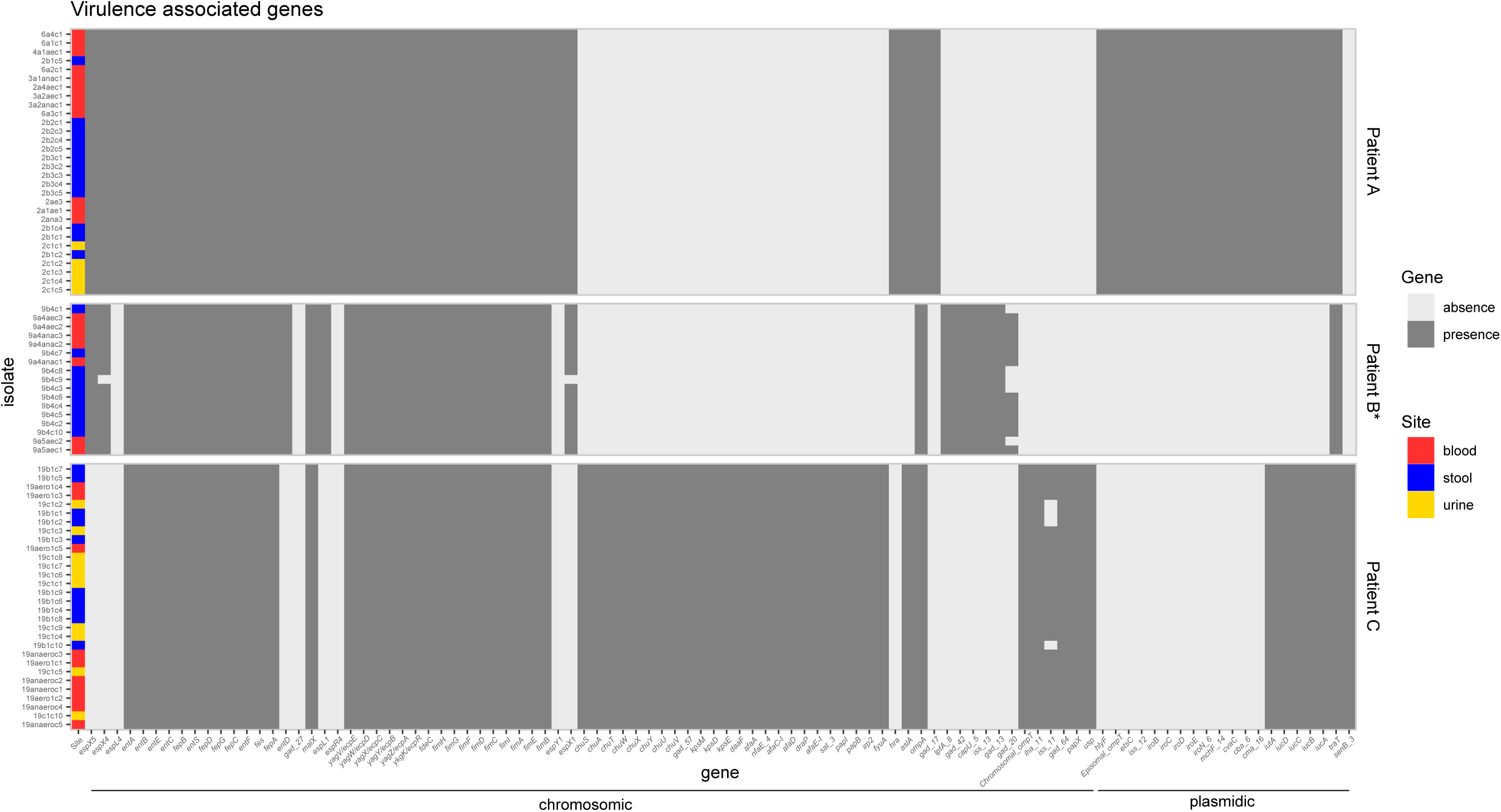
Presence/absence heatmaps of virulence associated genes of *E. coli* isolates when compared to the pan-virulome (including the virulence genes of all isolates). We considered a gene as present when at least 80% of its length was covered by at least one read. Genes are ordered by synteny on contigs. All isolates of patient B are mutators (*). The prevailing predicted localization of genes by PlaScope (chromosomic or plasmidic) is indicated (full list in supplementary Table S9). Note that plasmidic genes are not mobile, at the opposite of resistance genes (see Figure 5).

As a control for false-negative results, we evaluated the presence of 14 MLST genes and of genes encoded on plasmids. We recovered all the MLST genes for all patients (**Figure S3).** With a more conservative threshold (at least 5 reads covering more than 80% of the gene), we also found all the MLST genes for patient A and C. For patient B (mutator strain), 6 genes were not recovered over the 238 possibilities (**Figure S3**). Regarding genes encoded on a plasmid, there were very few differences depending on the depth threshold used (**Figure S4-S5**). With a more stringent threshold, we found 6 additional missing genes, over 4710 possibilities, for patient A and 7 additional missing genes, over 6477 possibilities for patient B. We did not find any differences for patient C. For patient A and B, in most of the cases missing genes corresponded to the absence of all the genes encoded on the corresponding plasmid. There were few exceptions that could be explained by a recent loss of the plasmid either *in vivo*, when isolates of the same cluster lack the same plasmid (*e.g.* plasmid 2 of patient A), or *in vitro* during the lab subcultures as we detected traces of the lost plasmids (*i.e.* portions of some of the genes encoded on the plasmid) suggesting the presence of the plasmid at a low frequency in the sequenced colony.

### Genomic traces of selection

We computed dN/dS ratios to assess whether gene sequences evolved neutrally or were under purifying or diversifying selection (**Figure 7, Table S10**). As expected, all isolates of patient B were under neutral selection (67, 68). For patients A and C, the same pattern of selection was found for blood and stool isolates, diversifying selection for patient A and purifying selection for patient C. We also found that all isolates sampled in urine evolved under neutral selection (patients A and C). As such results might be biased when the phylogenetic history is likely to include changes that occurred in other compartments, we computed dN/dS ratios in clades grouping samples of the same compartment (stool or blood). Despite the limited number of samples preventing us from obtaining significant results, we globally found the same trends in selection patterns (**Table S11**).

**Figure 7.**
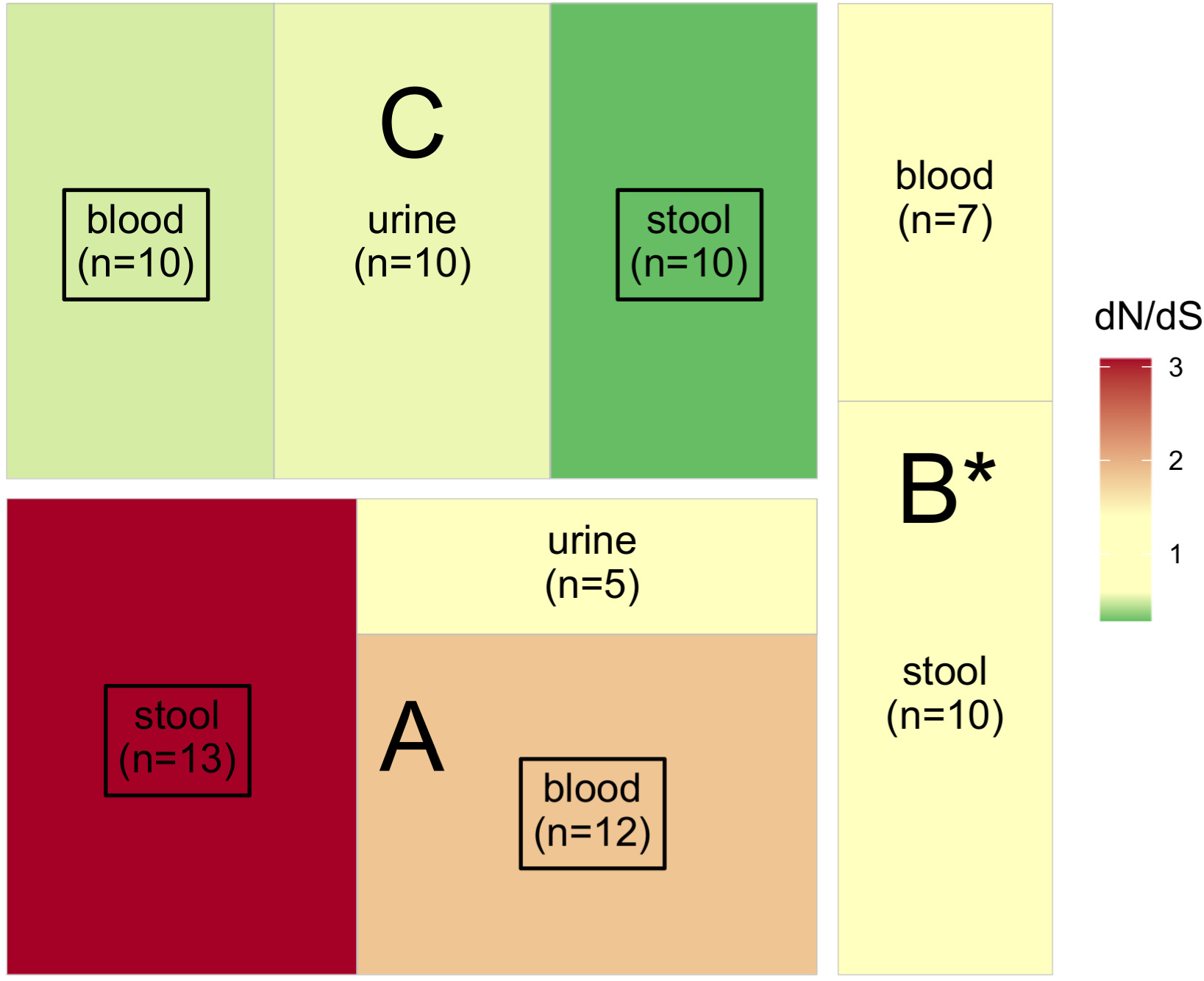
Action of selection on sequences (dN/dS) of *E. coli* isolates for patients A, B and C. Significant results are framed in black (see Table S9). All isolates of patient B are mutators (*). Neutrality (dN/dS not significantly different from 0) is indicated in pale yellow whereas purifying and diversifying selection are indicated in green and red, respectively.

This analysis does not compute the variation in non-coding regions; however, it encompasses more than 80% of all the variants and more than 87% of the genome. It should be noted that the number of variants (missense and synonymous) cannot be directly compared to dN/dS ratios (**Table 1**). These variants were computed with reference to a single sequence whereas ratios were computed against a reconstructed ancestral sequence.

## Discussion

The bacterial species *E. coli* is an opportunistic pathogen (6, 9) which may cross the intestinal barrier to reach extra-intestinal compartments causing infections. In this context, it has been proposed that virulence could be a by-product of commensalism (8). While evidence for purifying and neutral selection has been detected in commensal *E. coli* (10–12), extra-intestinal infection isolates have been shown to be under strong adaptive selection (13). Here we investigated both commensal and extra-intestinal isolates from leukemia patients undergoing chemotherapy. We observed the same strain in all compartments. Isolates of patients A and C were characterized by a very limited number of SNPs, 5 in average (ranging from 0 to 14) (69) and patient B was infected by a mutator strain (64). Interestingly, while each patient had a different selection signature, it was identical in both the bloodstream and gut. In urine, in all cases, neutral selection was at play.

### Gut origin of urine and blood immunosuppressed patient isolates

For each patient, the isolates collected from the gut, urine, and blood displayed highly similar sequences. They share the same phylogroup, ST, serotype, and *fimH* allele (**Table 1**), indicating that the strain most likely originated from the gut, as previously shown (31, 34). Moreover, with the exception of the mutator isolates, there were fewer than 15 SNPs between pairs of isolates of the same patient (**Figure 2**), the resistance and virulence profiles were stable and there were few differences in the IS repertoire (**Figures 5-6**). The phylogenetic distribution of isolates, together with the SNPs and IS distribution among compartments, suggested the translocation of the gut population diversity to extra-intestinal compartments for all patients (**Figures 3-4**).

### Same adaptation forces at play in the gut and bloodstream of immunosuppressed patient

In non-immunosuppressed patients, purifying (or neutral) evolution of *E. coli* populations has been reported in the gut (12) but not in extra-intestinal compartments where bacteria DNA sequence are under diversifying selection with strong evidence for gene level convergence (13).

Here, we found various signatures of selection in gut isolates. However, the same adaptive process was at play in the bloodstream and in the gut for each patient. For patient B, the presence of mutators at high-frequency in the population (all isolates) is indirect evidence for ongoing selection (70, 71). Indeed, even if they will ultimately be counter-selected (64), mutator alleles can reach 100% frequency in a population because they are associated with beneficial mutations (genetic hitchhiking) (72). In natural populations, mutators are present up to 15% frequency (73– 76). Despite the low number of patients, we noted a higher frequency (33%), that might be explained by patient treatment. Indeed, some anticancer chemotherapy drugs enhance the bacterial mutagenesis, thus promoting the emergence of a mutator clone (77). The presence of mutators at high frequency suggests a recent adaptative episode with a temporary indirect selection of this phenotype. Then, the mutator phenotype, even if the high mutation load observed might already have reduced the strain fitness, did not inhibit this *E. coli* population from causing a bloodstream infection. In patient A, diversifying selection is at play, whereas purifying selection shaped the isolates of patient C. Differences in the microbiota perturbation of these patients undergoing a specific antibiotic therapy and chemotherapy could explain this discrepancy. Unlike in extra-intestinal infection of non-immunosuppressed patients (13), no convergence among patients was detected. Moreover, isolates were almost identical with a very low number of SNPs differentiating them (five on average) unspecific to the compartment. Whereas, in non-immunosuppressed patients, niche adaptation was evidenced by specific mutations associated with isolates from the gut or the bladder (28). A weakened immune system and a permeable intestinal barrier, due to chemotherapy and antibiotic treatments, likely contribute to this this phenomenon.

### Neutral selection in the bladder of immunosuppressed patient

For both patients with urine samples, isolates evolved neutrally. Fluctuating environment or a small population size could explain the weak strength of selection observed in the commensal habitat of *E. coli*. Indeed, bacteria in the bladder face continuous adaptive challenges including fluctuations in exposure to the host immune system, antibiotic treatments, nutrients availability and a diverse microbial community due to an alternation between storage and voiding phases (78).

Our work obviously presents a major limitation as we evaluated a very limited number of patients, which prevents us from overgeneralizing our conclusions. Collecting stool before antibiotic treatment is indeed very difficult to achieve in clinical practices. Nevertheless, the evaluation of larger cohorts is necessary to associate evolution patterns and the environment (patient follow-up and patient-dependent factors) and to decipher the factors linked to the selection patterns observed here.

## Conclusion

We evaluated the diversity of *E. coli* isolates in stool, blood and urine of immunocompromised patients and found the same strain across compartments. We showed that all the diversity of the population is translocating. However, contrary to non-immunocompromised patients, we did not detect any modifications in the adaptive constraints between the gut and the bloodstream. Such multi-site sampling studies should be performed in non-immunosuppressed patients to strengthen our findings.

## Data availability

The data generated in this study have been submitted to the NCBI BioProject database under the accession number PRJEB69525 [https://www.ncbi.nlm.nih.gov/bioproject/?term=PRJEB69525].

## Acknowledgments

We are grateful to the INRAE MIGALE bioinformatics facility (MIGALE, INRAE, 2020. Migale bioinformatics Facility, doi: 10.15454/1.5572390655343293E12) for providing computing resources. We would also like to thank the Molecular Biology and Data Processing Platform of the Sorbonne Paris Nord University. This study was partly financed by two grants: one awarded by the research commission of the Université Sorbonne Paris Nord (BQR) and the other by the IFRB, UFR SMBH, Université Sorbonne Paris Nord.

## References

1. Pagano L, Caira M, Rossi G, Tumbarello M, Fanci R, Garzia MG, Vianelli N, Filardi N, De Fabritiis P, Beltrame A, Musso M, Piccin A, Cuneo A, Cattaneo C, Aloisi T, Riva M, Rossi G, Salvadori U, Brugiatelli M, Sannicolò S, Morselli M, Bonini A, Viale P, Nosari A, Aversa F, for the Hema e-Chart Group I. 2012. A prospective survey of febrile events in hematological malignancies. Ann Hematol 91:767–774.

2. Klastersky J, Ameye L, Maertens J, Georgala A, Muanza F, Aoun M, Ferrant A, Rapoport B, Rolston K, Paesmans M. 2007. Bacteraemia in febrile neutropenic cancer patients. International Journal of Antimicrobial Agents 30:51–59.

3. Tumbarello M, Spanu T, Caira M, Trecarichi EM, Laurenti L, Montuori E, Fianchi L, Leone F, Fadda G, Cauda R, Pagano L. 2009. Factors associated with mortality in bacteremic patients with hematologic malignancies. Diagnostic Microbiology and Infectious Disease 64:320–326.

4. Trecarichi EM, Pagano L, Candoni A, Pastore D, Cattaneo C, Fanci R, Nosari A, Caira M, Spadea A, Busca A, Vianelli N, Tumbarello M. 2015. Current epidemiology and antimicrobial resistance data for bacterial bloodstream infections in patients with hematologic malignancies: an Italian multicentre prospective survey. Clinical Microbiology and Infection 21:337–343.

5. Montassier E, Batard E, Gastinne T, Potel G, de La Cochetière MF. 2013. Recent changes in bacteremia in patients with cancer: a systematic review of epidemiology and antibiotic resistance. Eur J Clin Microbiol Infect Dis 32:841–850.

6. Tenaillon O, Skurnik D, Picard B, Denamur E. 2010. The population genetics of commensal Escherichia coli. Nat Rev Microbiol 8:207–217.

7. Diard M, Garry L, Selva M, Mosser T, Denamur E, Matic I. 2010. Pathogenicity-Associated Islands in Extraintestinal Pathogenic Escherichia coli Are Fitness Elements Involved in Intestinal Colonization. Journal of Bacteriology 192:4885–4893.

8. Le Gall T, Clermont O, Gouriou S, Picard B, Nassif X, Denamur E, Tenaillon O. 2007. Extraintestinal Virulence Is a Coincidental By-Product of Commensalism in B2 Phylogenetic Group Escherichia coli Strains. Molecular Biology and Evolution 24:2373–2384.

9. Denamur E, Clermont O, Bonacorsi S, Gordon D. 2021. The population genetics of pathogenic Escherichia coli. Nat Rev Microbiol 19:37–54.

10. Schloissnig S, Arumugam M, Sunagawa S, Mitreva M, Tap J, Zhu A, Waller A, Mende DR, Kultima JR, Martin J, Kota K, Sunyaev SR, Weinstock GM, Bork P. 2013. Genomic variation landscape of the human gut microbiome. Nature 493:45–50.

11. Hu D, Fuller NR, Caterson ID, Holmes AJ, Reeves PR. 2022. Single-gene long-read sequencing illuminates Escherichia coli strain dynamics in the human intestinal microbiome. Cell Rep 38:110239.

12. Ghalayini M, Launay A, Bridier-Nahmias A, Clermont O, Denamur E, Lescat M, Tenaillon O. 2018. Evolution of a Dominant Natural Isolate of Escherichia coli in the Human Gut over the Course of a Year Suggests a Neutral Evolution with Reduced Effective Population Size. Applied and Environmental Microbiology 84:e02377–17.

13. Bridier-Nahmias A, Launay A, Bleibtreu A, Magnan M, Walewski V, Chatel J, Dion S, Robbe-Saule V, Clermont O, Norel F, Denamur E, Tenaillon O. 2021. Escherichia coli Genomic Diversity within Extraintestinal Acute Infections Argues for Adaptive Evolution at Play. mSphere 6:e01176–20.

14. Lieberman TD, Michel J-B, Aingaran M, Potter-Bynoe G, Roux D, Davis MR, Skurnik D, Leiby N, LiPuma JJ, Goldberg JB, McAdam AJ, Priebe GP, Kishony R. 2011. Parallel bacterial evolution within multiple patients identifies candidate pathogenicity genes. 12. Nat Genet 43:1275–1280.

15. Yang L, Jelsbak L, Marvig RL, Damkiær S, Workman CT, Rau MH, Hansen SK, Folkesson A, Johansen HK, Ciofu O, Høiby N, Sommer MOA, Molin S. 2011. Evolutionary dynamics of bacteria in a human host environment. Proceedings of the National Academy of Sciences 108:7481–7486.

16. Marvig RL, Sommer LM, Molin S, Johansen HK. 2015. Convergent evolution and adaptation of Pseudomonas aeruginosa within patients with cystic fibrosis. 1. Nat Genet 47:57–64.

17. Jorth P, Staudinger BJ, Wu X, Hisert KB, Hayden H, Garudathri J, Harding CL, Radey MC, Rezayat A, Bautista G, Berrington WR, Goddard AF, Zheng C, Angermeyer A, Brittnacher MJ, Kitzman J, Shendure J, Fligner CL, Mittler J, Aitken ML, Manoil C, Bruce JE, Yahr TL, Singh PK. 2015. Regional Isolation Drives Bacterial Diversification within Cystic Fibrosis Lungs. Cell Host & Microbe 18:307–319.

18. Ashish A, Paterson S, Mowat E, Fothergill JL, Walshaw MJ, Winstanley C. 2013. Extensive diversification is a common feature of Pseudomonas aeruginosa populations during respiratory infections in cystic fibrosis. Journal of Cystic Fibrosis 12:790–793.

19. Darch SE, McNally A, Harrison F, Corander J, Barr HL, Paszkiewicz K, Holden S, Fogarty A, Crusz SA, Diggle SP. 2015. Recombination is a key driver of genomic and phenotypic diversity in a Pseudomonas aeruginosa population during cystic fibrosis infection. 1. Sci Rep 5:7649.

20. Lieberman TD, Flett KB, Yelin I, Martin TR, McAdam AJ, Priebe GP, Kishony R. 2014. Genetic variation of a bacterial pathogen within individuals with cystic fibrosis provides a record of selective pressures. 1. Nat Genet 46:82–87.

21. Williams D, Evans B, Haldenby S, Walshaw MJ, Brockhurst MA, Winstanley C, Paterson S. 2015. Divergent, Coexisting Pseudomonas aeruginosa Lineages in Chronic Cystic Fibrosis Lung Infections. Am J Respir Crit Care Med 191:775–785.

22. Suerbaum S, Josenhans C. 2007. Helicobacter pylori evolution and phenotypic diversification in a changing host. 6. Nat Rev Microbiol 5:441–452.

23. Smith EE, Buckley DG, Wu Z, Saenphimmachak C, Hoffman LR, D’Argenio DA, Miller SI, Ramsey BW, Speert DP, Moskowitz SM, Burns JL, Kaul R, Olson MV. 2006. Genetic adaptation by Pseudomonas aeruginosa to the airways of cystic fibrosis patients. Proceedings of the National Academy of Sciences 103:8487–8492.

24. Moxon ER, Rainey PB, Nowak MA, Lenski RE. 1994. Adaptive evolution of highly mutable loci in pathogenic bacteria. Current Biology 4:24–33.

25. van der Woude MW, Bäumler AJ. 2004. Phase and Antigenic Variation in Bacteria. Clinical Microbiology Reviews 17:581–611.

26. Musher DM, Dowell ME, Shortridge VD, Flamm RK, Jorgensen JH, Le Magueres P, Krause KL. 2002. Emergence of Macrolide Resistance during Treatment of Pneumococcal Pneumonia. New England Journal of Medicine 346:630–631.

27. Wong A, Kassen R. 2011. Parallel evolution and local differentiation in quinolone resistance in Pseudomonas aeruginosa. Microbiology 157:937–944.

28. Thänert R, Choi J, Reske KA, Hink T, Thänert A, Wallace MA, Wang B, Seiler S, Cass C, Bost MH, Struttmann EL, Iqbal ZH, Sax SR, Fraser VJ, Baker AW, Foy KR, Williams B, Xu B, Capocci-Tolomeo P, Lautenbach E, Burnham C-AD, Dubberke ER, Kwon JH, Dantas G. 2022. Persisting uropathogenic Escherichia coli lineages show signatures of niche-specific within-host adaptation mediated by mobile genetic elements. Cell Host & Microbe 30:1034–1047.e6.

29. Kessler R, Nisa S, Hazen TH, Horneman A, Amoroso A, Rasko DA, Donnenberg MS. 2015. Diarrhea, bacteremia and multiorgan dysfunction due to an extraintestinal pathogenic Escherichia coli strain with enteropathogenic E. coli genes. Pathogens and Disease 73.

30. Maslow JN, Mulligan ME, Arbeit RD. 1994. Recurrent Escherichia coli bacteremia. Journal of Clinical Microbiology 32:710–714.

31. Berg RD. 1995. Bacterial translocation from the gastrointestinal tract. Trends in Microbiology 3:149–154.

32. van Vliet MJ, Tissing WJE, Dun CAJ, Meessen NEL, Kamps WA, de Bont ESJM, Harmsen HJM. 2009. Chemotherapy Treatment in Pediatric Patients with Acute Myeloid Leukemia Receiving Antimicrobial Prophylaxis Leads to a Relative Increase of Colonization with Potentially Pathogenic Bacteria in the Gut. Clinical Infectious Diseases 49:262–270.

33. van Vliet MJ, Harmsen HJM, Bont ESJM de, Tissing WJE. 2010. The Role of Intestinal Microbiota in the Development and Severity of Chemotherapy-Induced Mucositis. PLOS Pathogens 6:e1000879.

34. Samet A, Śledzińska A, Krawczyk B, Hellmann A, Nowicki S, Kur J, Nowicki B. 2013. Leukemia and risk of recurrent Escherichia coli bacteremia: genotyping implicates E. coli translocation from the colon to the bloodstream. Eur J Clin Microbiol Infect Dis 32:1393–1400.

35. Clermont O, Christenson JK, Denamur E, Gordon DM. 2013. The Clermont Escherichia coli phylo-typing method revisited: improvement of specificity and detection of new phylo-groups. Environmental Microbiology Reports 5:58–65.

36. Bankevich A, Nurk S, Antipov D, Gurevich AA, Dvorkin M, Kulikov AS, Lesin VM, Nikolenko SI, Pham S, Prjibelski AD, Pyshkin AV, Sirotkin AV, Vyahhi N, Tesler G, Alekseyev MA, Pevzner PA. 2012. SPAdes: A New Genome Assembly Algorithm and Its Applications to Single-Cell Sequencing. Journal of Computational Biology 19:455–477.

37. Wick R. 2022. rrwick/Filtlong. C++.

38. Wick RR, Judd LM, Gorrie CL, Holt KE. 2017. Unicycler: Resolving bacterial genome assemblies from short and long sequencing reads. PLOS Computational Biology 13:e1005595.

39. Seemann T. 2014. Prokka: rapid prokaryotic genome annotation. Bioinformatics 30:2068– 2069.

40. Royer G, Decousser JW, Branger C, Dubois M, Médigue C, Denamur E, Vallenet D. 2018. PlaScope: a targeted approach to assess the plasmidome from genome assemblies at the species level. Microbial Genomics 4:e000211.

41. Page AJ, Cummins CA, Hunt M, Wong VK, Reuter S, Holden MTG, Fookes M, Falush D, Keane JA, Parkhill J. 2015. Roary: rapid large-scale prokaryote pan genome analysis. Bioinformatics 31:3691–3693.

42. Martin M. 2011. Cutadapt removes adapter sequences from high-throughput sequencing reads. 1. EMBnet.journal 17:10–12.

43. Seemann T. 2015. Snippy: fast bacterial variant calling from NGS reads.

44. Li H. 2013. Aligning sequence reads, clone sequences and assembly contigs with BWA-MEM. arXiv:13033997 [q-bio].

45. Kronenberg ZN, Osborne EJ, Cone KR, Kennedy BJ, Domyan ET, Shapiro MD, Elde NC, Yandell M. 2015. Wham: Identifying Structural Variants of Biological Consequence. PLOS Computational Biology 11:e1004572.

46. Siguier P, Perochon J, Lestrade L, Mahillon J, Chandler M. 2006. ISfinder: the reference centre for bacterial insertion sequences. Nucleic Acids Research 34:D32–D36.

47. Treepong P, Guyeux C, Meunier A, Couchoud C, Hocquet D, Valot B. 2018. *panISa: ab initio* detection of insertion sequences in bacterial genomes from short read sequence data. Bioinformatics 34:3795–3800.

48. Bourrel AS, Poirel L, Royer G, Darty M, Vuillemin X, Kieffer N, Clermont O, Denamur E, Nordmann P, Decousser J-W, IAME Resistance Group. 2019. Colistin resistance in Parisian inpatient faecal Escherichia coli as the result of two distinct evolutionary pathways. Journal of Antimicrobial Chemotherapy 74:1521–1530.

49. Inouye M, Dashnow H, Raven L-A, Schultz MB, Pope BJ, Tomita T, Zobel J, Holt KE. 2014. SRST2: Rapid genomic surveillance for public health and hospital microbiology labs. Genome Medicine 6:90.

50. Jaureguy F, Landraud L, Passet V, Diancourt L, Frapy E, Guigon G, Carbonnelle E, Lortholary O, Clermont O, Denamur E, Picard B, Nassif X, Brisse S. 2008. Phylogenetic and genomic diversity of human bacteremic Escherichia coli strains. BMC Genomics 9:560.

51. Wirth T, Falush D, Lan R, Colles F, Mensa P, Wieler LH, Karch H, Reeves PR, Maiden MCJ, Ochman H, Achtman M. 2006. Sex and virulence in Escherichia coli: an evolutionary perspective. Molecular Microbiology 60:1136–1151.

52. Ingle DJ, Valcanis M, Kuzevski A, Tauschek M, Inouye M, Stinear T, Levine MM, Robins-Browne RM, Holt KE. 2016. In silico serotyping of E. coli from short read data identifies limited novel O-loci but extensive diversity of O:H serotype combinations within and between pathogenic lineages. Microbial Genomics 2.

53. Roer L, Tchesnokova V, Allesøe R, Muradova M, Chattopadhyay S, Ahrenfeldt J, Thomsen MCF, Lund O, Hansen F, Hammerum AM, Sokurenko E, Hasman H. 2017. Development of a Web Tool for Escherichia coli Subtyping Based on fimH Alleles. Journal of Clinical Microbiology 55:2538–2543.

54. Beghain J, Bridier-Nahmias A, Le Nagard H, Denamur E, Clermont O. 2018. ClermonTyping: an easy-to-use and accurate in silico method for Escherichia genus strain phylotyping. Microb Genom 4:e000192.

55. Zankari E, Hasman H, Cosentino S, Vestergaard M, Rasmussen S, Lund O, Aarestrup FM, Larsen MV. 2012. Identification of acquired antimicrobial resistance genes. Journal of Antimicrobial Chemotherapy 67:2640–2644.

56. Chen L, Zheng D, Liu B, Yang J, Jin Q. 2016. VFDB 2016: hierarchical and refined dataset for big data analysis—10 years on. Nucleic Acids Research 44:D694–D697.

57. Joensen KG, Scheutz F, Lund O, Hasman H, Kaas RS, Nielsen EM, Aarestrup FM. 2014. Real-Time Whole-Genome Sequencing for Routine Typing, Surveillance, and Outbreak Detection of Verotoxigenic Escherichia coli. Journal of Clinical Microbiology 52:1501–1510.

58. Marin J, Clermont O, Royer G, Mercier-Darty M, Decousser JW, Tenaillon O, Denamur E, Blanquart F. 2022. The Population Genomics of Increased Virulence and Antibiotic Resistance in Human Commensal Escherichia coli over 30 Years in France. Applied and Environmental Microbiology 0:e00664–22.

59. R Developement Core Team. 2010. R: A language and environment for statistical computing. (No Title).

60. Didelot X, Croucher NJ, Bentley SD, Harris SR, Wilson DJ. 2018. Bayesian inference of ancestral dates on bacterial phylogenetic trees. Nucleic Acids Research 46:e134.

61. Schliep KP. 2011. phangorn: phylogenetic analysis in R. Bioinformatics 27:592–593.

62. Manning CD, Raghavan P, Schütze H. 2008. Introduction to information retrieval. Cambridge University Press, New York.

63. Oliver A, Cantón R, Campo P, Baquero F, Blázquez J. 2000. High Frequency of Hypermutable Pseudomonas aeruginosa in Cystic Fibrosis Lung Infection. Science 288:1251–1253.

64. Touchon M, Perrin A, Sousa JAM de, Vangchhia B, Burn S, O’Brien CL, Denamur E, Gordon D, Rocha EP. 2020. Phylogenetic background and habitat drive the genetic diversification of Escherichia coli. PLOS Genetics 16:e1008866.

65. Fehér T, Bogos B, Méhi O, Fekete G, Csörgő B, Kovács K, Pósfai G, Papp B, Hurst LD, Pál C. 2012. Competition between Transposable Elements and Mutator Genes in Bacteria. Molecular Biology and Evolution 29:3153–3159.

66. Maharjan RP, Liu B, Li Y, Reeves PR, Wang L, Ferenci T. 2013. Mutation accumulation and fitness in mutator subpopulations of Escherichia coli. Biology Letters 9:20120961.

67. Kishimoto T, Iijima L, Tatsumi M, Ono N, Oyake A, Hashimoto T, Matsuo M, Okubo M, Suzuki S, Mori K, Kashiwagi A, Furusawa C, Ying B-W, Yomo T. 2010. Transition from Positive to Neutral in Mutation Fixation along with Continuing Rising Fitness in Thermal Adaptive Evolution. PLoS Genet 6:e1001164.

68. Duval A, Opatowski L, Brisse S. 2023. Defining genomic epidemiology thresholds for common-source bacterial outbreaks: a modelling study. The Lancet Microbe 4:e349–e357.

69. Denamur E, Matic I. 2006. Evolution of mutation rates in bacteria. Molecular Microbiology 60:820–827.

70. Tenaillon O, Toupance B, Le Nagard H, Taddei F, Godelle B. 1999. Mutators, Population Size, Adaptive Landscape and the Adaptation of Asexual Populations of Bacteria. Genetics 152:485–493.

71. Sniegowski PD, Gerrish PJ, Lenski RE. 1997. Evolution of high mutation rates in experimental populations of E. coli. 6634. Nature 387:703–705.

72. Smith JM, Haigh J. 1974. The hitch-hiking effect of a favourable gene. Genetics Research 23:23–35.

73. Jyssum K. 1960. Observations on Two Types of Genetic Instability in Escherichia Coli. Acta Pathologica Microbiologica Scandinavica 48:113–120.

74. Gross MD, Siegel EC. 1981. Incidence of mutator strains in Escherichia coli and coliforms in nature. Mutation Research Letters 91:107–110.

75. LeClerc JE, Li B, Payne WL, Cebula TA. 1996. High Mutation Frequencies Among Escherichia coli and Salmonella Pathogens. Science 274:1208–1211.

76. Matic I, Radman M, Taddei F, Picard B, Doit C, Bingen E, Denamur E, Elion J. 1997. Highly Variable Mutation Rates in Commensal and Pathogenic Escherichia coli. Science 277:1833– 1834.

77. Hobson CA, Bonacorsi S, Hocquet D, Baruchel A, Fahd M, Storme T, Tang R, Doit C, Tenaillon O, Birgy A. 2020. Impact of anticancer chemotherapy on the extension of beta-lactamase spectrum: an example with KPC-type carbapenemase activity towards ceftazidime-avibactam. 1. Sci Rep 10:589.

78. Vahabi B, Drake MJ. 2015. Physiological and pathophysiological implications of micromotion activity in urinary bladder function. Acta Physiologica 213:360–370.

